# Membrane potential phase shifts differ for excitation vs. inhibition in resonant pyramidal neurons: a computer modeling study

**DOI:** 10.1101/2023.03.20.533519

**Authors:** Craig Kelley, Srdjan D. Antic, Nicholas T. Carnevale, John L. Kubie, William W. Lytton

**Affiliations:** Program in Biomedical Engineering, SUNY Downstate Health Sciences University & NYU Tandon School of Engineering, Brooklyn, NY, USA; Institute of Systems Genomics, Neuroscience Department, University of Connecticut Health, Farmington, CT, USA; Department of Neuroscience, Yale University, New Haven, CT, USA; The Robert F. Furchgott Center for Neural and Behavioral Science, Brooklyn, NY, USA; Department of Cell Biology, SUNY Downstate Health Sciences University, Brooklyn, NY, USA2; Department of Physiology & Pharmacology, SUNY Downstate Health Sciences University, Brooklyn, NY, USA; Department of Neurology, SUNY Downstate Health Sciences University, Brooklyn, NY, USA; Department of Neurology, Kings County Hospital Center, Brooklyn, NY, USA; Aligning Science Across Parkinson’s (ASAP) Collaborative Research Network, Chevy Chase, MD USA

## Abstract

Rhythmic activity is ubiquitous in neural systems, and impedance analysis has been widely used to examine frequency-dependent responses of neuronal membranes to rhythmic inputs. Impedance analysis assumes the neuronal membrane is a linear system, requiring the use of small signals to stay in a near-linear regime. However, postsynaptic potentials are often large and trigger nonlinear mechanisms. We therefore augmented impedance analysis to evaluate membrane responses in this nonlinear domain, analyzing responses to injected current for subthreshold membrane voltage (V_memb_), suprathreshold spike-blocked V_memb_, and spiking in a validated neocortical pyramidal neuron computer model. Responses in these output regimes were asymmetrical, with different phase shifts during hyperpolarizing and depolarizing half-cycles. Suprathreshold chirp stimulation gave equivocal results due to nonstationarity of response, requiring us to use fixed-frequency sinusoids. Sinusoidal inputs produced *phase retreat*: action potentials occurred progressively later in cycles of the input stimulus, resulting from adaptation. Conversely, sinusoidal current with increasing amplitude over cycles produced a pattern of *phase advance*: action potentials occurred progressively earlier. Phase retreat was dependent on *I*_h_ and *I*_AHP_ currents; phase advance was modulated by these currents. Our results suggest differential responses of cortical neurons depending on the frequency of oscillatory input in the delta – beta range, which will play a role in neuronal responses to shifts in network state. We hypothesize that intrinsic cellular properties complement network properties and contribute to *in vivo* phase-shift phenomena such as phase precession, seen in place and grid cells, and phase roll, observed in hippocampal CA1 neurons.

**New & Noteworthy:** We augmented electrical impedance analysis to characterize phase shifts between large amplitude current stimuli and nonlinear, asymmetric membrane potential responses. We predict different frequency-dependent phase shifts in response excitation versus inhibition, as well as shifts in spike timing over multiple input cycles, in resonant pyramidal neurons. We hypothesize that these effects contribute to navigation-related phenomena like phase precession and phase roll. Our neuron-level hypothesis complements, rather than falsifies, prior network-level explanations of these phenomena.

## Introduction

Rhythmic activity is ubiquitous in cortical structures of the brain and can be measured as oscillations in the local field potentials (LFPs) recorded extracellularly. LFPs are largely generated by transmembrane currents, principally postsynaptic dendritic currents, in neurons near the recording electrode (Engel et al., 2001; Buzsáki et al., 2012), and recent work has suggested that pyramidal neurons are the largest contributors to these rhythms (Dura-Bernal et al., 2022). Pyramidal neurons have diverse biophysics and projections, forming the main outputs of cortical circuits (Aronoff et al., 2010; Harris and Shepherd, 2015; Hattox and Nelson, 2007). Understanding how pyramidal neurons integrate oscillatory synaptic inputs to generate subthreshold membrane potential oscillations and spiking activity is therefore critical to understanding cortical computations (Engel et al., 2001; Fries et al., 2007; Harris et al., 2002; Mehta et al., 2002). Here we focus on the neocortical layer 5b (L5b) pyramidal neuron, which integrates inputs from thalamus, other cortical layers, and other cortical areas, and exerts top-down control over subcortical brain areas (Agmon and Connors, 1992; Meyer et al., 2010; Wimmer et al., 2010; Oberlaender et al., 2012; Rah et al., 2013; Markram et al., 2015). Neocortical L5b pyramidal neurons, like hippocampal CA1 pyramidal neurons and entorhinal pyramidal neurons, are theta-resonant (membrane potential – V_memb_– responds most strongly to subthreshold oscillatory inputs 3-12 Hz) (Hutcheon et al., 1996; Ulrich, 2002; Leung and Yu, 1998; Binini et al., 2021). Impedance analysis provides a tool for characterizing how the neuronal membrane responds to and filters oscillatory inputs.

The impedance analysis framework derives from electrical engineering and is based on linear circuits composed of resistors (R), capacitors (C), and inductors (L). Impedance analysis assumes that the neuronal membrane is a linear, time-invariant (stationary) system. In a neural context, linearity means that positive and negative current stimuli of equivalent amplitude will produce equal, opposite, and additive changes to V_memb_, which is often not the case. *Nonlinearity* of neuronal membrane response arises through voltage-gated ion channels, whose conductances are different during depolarization and hyperpolarization. Stationarity (time-invariance) implies that the membrane always has the same response to equivalent inputs. *Nonstationarity* arises from intrinsic mechanisms that preserve history (history dependence) – slow channel kinetics, Ca^2+^ accumulation, neuromodulation (changing state or set-points), *etc*. Classical impedance analysis is therefore ill-suited for studying neuronal responses, which are markedly nonlinear and time-varying (Miller et al., 1985; Rotstein, 2014). Nonetheless, classical impedance analysis has been widely used for signals in a low-amplitude regime where the primary nonlinearity is approximated by a linear inductance, considered as a “phenomenological inductance” (Cole, 1941; Mauro et al., 1970).

Neuronal membrane impedance is usually estimated by applying current as either a chirp (time-varying signal whose instantaneous frequency increases with time) (Puil et al., 1986) or white noise (Moore and Christensen, 1985; Wright et al., 1996) and recording the resulting changes in V_memb_. Impedance is expressed as a complex number from which we can extract amplitude (|*Z*| – the frequency-dependent output strength) and phase (Φ – the frequency-dependent temporal shifts between corresponding points such as the peak or the trough in the current stimulus vs. that of the V_memb_ waveform). Because these methods properly pertain to a non-physiological, linear, time-invariant/stationary system, we developed new methods for characterizing the nonlinear filtering properties of the neuronal membrane, whether spiking or subthreshold.

A number of techniques have been used to expand the impedance framework to physiological signaling, with a focus on examining amplitude nonlinearities. For example, theta-frequency inputs were demonstrated to have an *I*_h_-dependent nonlinearity in hippocampal pyramidal neurons (Das and Narayanan, 2017), based on spike-triggered averaging (Bryant and Segundo, 1976). Ulrich (2002) used chirp stimulation of neocortical L5b pyramidal neurons to produce “band-passed spiking:” action potentials were largely seen in response to frequencies around the cell’s subthreshold resonance frequencies of 4 – 8 Hz. Asymmetric responses to excitation vs. inhibition have also been noted (Schreiber et al., 2009; Tohidi and Nadim, 2009; Rotstein, 2015). Pena et al. (2019) extended the impedance amplitude framework to assess these asymmetries, identifying separate resonance frequencies for hyperpolarizing and depolarizing voltage responses. Similarly, Fischer et al. (2017) identified the frequency dependence of leech-neuron spiking in response to sinusoidal inputs. Due to this focus on amplitude, the temporal characteristics of nonlinear responses have been largely ignored.

In this paper, we develop and apply a method analogous to impedance phase analysis for nonlinear V_memb_ responses, both sub- and suprathreshold. We used an experimentally-validated neocortical L5b pyramidal neuron model (Hay et al., 2011) whose location-dependent subthreshold impedance profile has also been validated (Kelley et al., 2021). We found nonlinear subthreshold V_memb_ responses to chirp stimuli maintained different frequency-dependent phase relationships during depolarization and hyperpolarization. However, chirp methods only presented an incomplete picture of these relationships due to nonstationarity of the V_memb_ response. By instead using continuous sinusoidal currents, we found that action potentials occurred progressively later in the cycles of the input stimulus, which we termed *phase retreat*. Conversely, increasing stimulus amplitude on each cycle produced progressively earlier action potentials, which we termed *phase advance*. We suggest that these processes are involved in relationships between local field potential oscillations and spike timing observed in a number of brain structures *in vivo* and often ascribed to network phenomena (Geisler et al., 2007; Jensen and Lisman, 1996; Bose and Recce, 2001; Wallenstein and Hasselmo, 1997; Tsodyks et al., 1996). Phase advance may contribute to phase precession, seen in place cells (O’Keefe and Recce, 1993), grid cells (Hafting et al., 2008), medial prefrontal cortex neurons (Jones and Wilson, 2005), and neurons in ventral striatum (van der Meer and Redish, 2011), and phase retreat may contribute to phase roll hippocampal CA1 neurons (Sloin et al., 2022).

## Methods

### Cell Model

In this study, we used a previous model of a neocortical layer 5b (L5b) pyramidal neuron (Hay et al., 2011; Kelley et al., 2021) – a thick-tufted (corticospinal, pyramidal-tract-type, PT) principle cell which projects to subcortical brain structures. The model incorporates a combination of *I*_h_ and TASK-like shunting current as described in Migliore and Migliore (2012), with constant HCN channel density in the soma and basal dendrites and exponentially increasing density with distance from the soma in apical dendrites. Using this model, we previously showed location-dependent, subthreshold impedance amplitude and phase profiles in close agreement with experimental observations (Kelley et al., 2021). Additional details of the original cell model can be found in Hay et al. (2011).

### Simulations & Analysis

As previously (Puil et al., 1986), a linear chirp current stimulus was defined as:

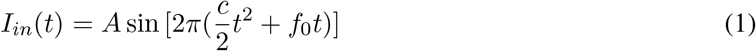

where *c* = (*f*_1_-*f*_0_) / *T, f*_0_ is the initial frequency in Hz, *f*_1_ is the final frequency, and *T* is the duration of the frequency sweep in seconds (*f*_1_ = 0.5; *f*_2_ = 12; *T* = 12). The stimulus amplitude, *A*, was chosen to be either subthreshold linear, subthreshold nonlinear, or suprathreshold with 1 spike/cycle at low frequency (≤12 Hz).

Since impedance analysis is inadequate for asymmetric, nonlinear responses, we extended a framework for computing a nonlinear analog of impedance amplitude developed by Pena et al. (2019) to compute a nonlinear analog of impedance phase (Φ) for depolarization and hyperpolarization. We defined the upper (+) and lower (-) envelopes of V_memb_ and *I*_*in*_ by finding the peaks 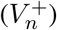 and troughs 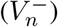 in V_memb_ and the peaks/troughs in the stimulus 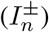 for each stimulus cycle *n* (Fig. 1). We defined the times of each peak/trough in V_memb_ as 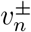 and the times of each peak/trough in the stimulus as 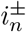. The instantaneous phase (*P* (*t*)) of the current stimulus was extracted using the Hilbert transform:

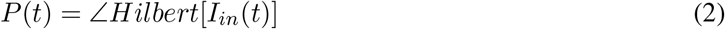

**Figure 1:**
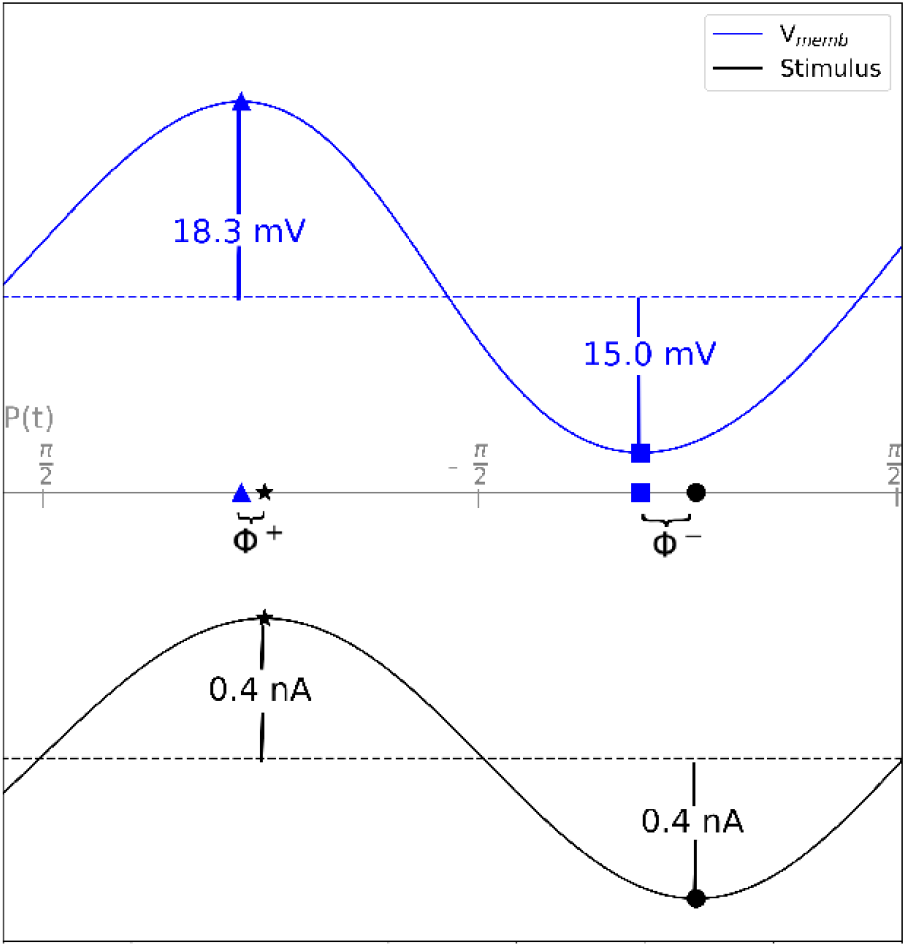
High amplitude stimuli produced nonlinear V_memb_ responses with different phase shifts for depolarization and hyperpolarization. Symmetrical, sinusoidal current stimulus (black, bottom) produced asymmetric V_memb_ response (blue, top). Superscripted plus (^+^) and minus (^−^) respectively label depolarizing and hyperpolarizing half-cycle peaks. Times of the V_memb_ peaks and troughs were mapped onto instantaneous stimulus phase. Difference in instantaneous stimulus phase between time of peak V_memb_ to peak in stimulus (triangle to star, Φ^+^) or trough to trough (square to circle Φ^−^) was the value of the phase shift: here, Φ^+^= 0.13, Φ^−^= 0.40 for a 3 Hz input. Both are positive, indicating phase lead (V_memb_ peak preceding stimulus peak) for both.

We defined our nonlinear Φ analog for the upper and lower V_memb_ envelopes (+/-, respectively) as:

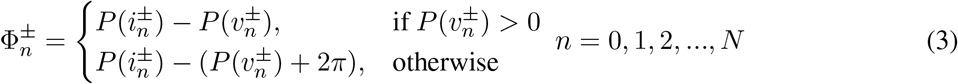

where *N* is the total number of stimulus cycles; the 2*π* correction was required when a V_memb_ trough *v*^−^ occurred after a stimulus trough *i*^−^, thus on the following cycle of *P* (*t*). Since the chirp stimulus has an instantaneous frequency that increased linearly with time, one can readily define 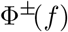, where *f* is stimulus frequency, analogously with 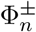. As with linear Φ, phase lead occurs when 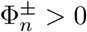: peaks/troughs of V_memb_ preceding peaks/troughs of *I*_*in*_(*t*). Similarly, phase lag is defined as 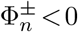, where peaks/troughs in V_memb_ lag behind the input stimulus. We defined a *synchrony point* as a time where 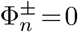, representing a transition point between lead and lag where peaks/troughs in V_memb_ and stimulus are synchronous. We computed 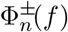 at the soma for chirp stimuli in subthreshold, suprathreshold, and spike-blockade suprathreshold conditions, comparing to the near-linear response to small subthreshold chirp, calculated using Fourier-based methods. For the suprathreshold spiking conditions, we computed Φ^+^ of a single spike or the first spike in a burst.

Because responses to suprathreshold chirp stimuli combined the effects of frequency shift with nonstationary cell response, we switched to regular sinusoidal stimuli to assess 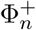 across cycles of constant frequency of 2-9 Hz. We also assessed sinusoidal stimuli of increasing amplitude: starting at 0 nA with linear increase of 0.32 nA/s unless noted otherwise.

In the course of this study, we ran more than 600 single-cell simulations. 1 second of simulation time required ∼10.5 seconds of clock-time in NEURON on a Linux system using a single core of a 1.60GHz Intel Core i5-10210U CPU. All simulations presented here were run using NEURON version 8.0 (Hines and Carnevale, 2001). The code developed for simulation, data analysis, and visualization was written in Python, and it is available on [GitHub](https://github.com/suny-downstate-medical-center/nonlinear_impedance_phase) and [ModelDB](https://modeldb.yale.edu/267651) (access code: heaviside).

## Results

Traditionally, impedance measures used for characterizing neuronal input/output (I/O; current/V_memb_) relationships relied on low-amplitude signals below the thresholds for activation of most voltage-sensitive ion channels (Mauro, 1961; Puil et al., 1986). This had the advantage of providing a near-linear I/O relationship that could be characterized using traditional methods. However, a major disadvantage was that it excluded larger, more physiologically-relevant signals with amplitudes comparable to compound postsynaptic potentials. In order to characterize the time course of neuronal responses to oscillatory stimuli in a more physiological regime, we provided higher amplitude inputs than typically used, high enough to activate voltage-sensitive ion channels. We then separated the phase measures for depolarizing peaks (Φ^+^) and hyperpolarizing troughs (Φ^−^), using the alternating half-cycles of the inputs (see *Methods*). These two phase measures were found to characterize very different I/O phase relationships, assessed both with and without spiking activity (Fig. 2). Evidence of nonstationarity in the phase relationship between spiking activity and stimulus (change in phase shift between spikes and stimulus over time), led us to assess responses to more prolonged fixed-frequency sinusoids in addition to chirps. Identified patterns demonstrated how the intrinsic electrophysiology of a single neuron could give rise to the phase shifts seen in behaving animals: phase precession (O’Keefe and Recce, 1993) and phase roll (Sloin et al., 2022).

**Figure 2:**
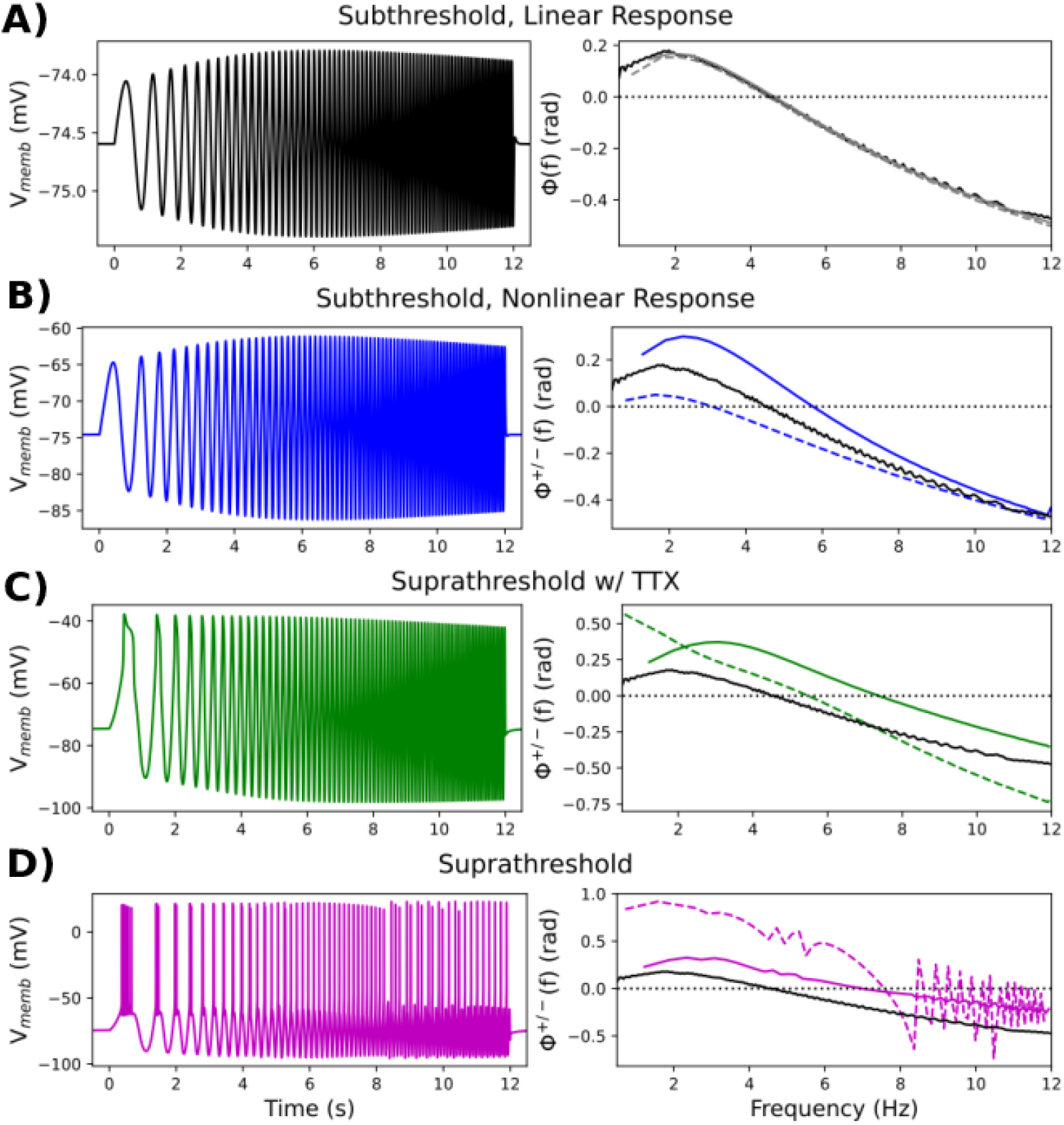
Higher amplitude chirp stimuli (input I) produced nonlinear V_memb_ responses (output O) with differing frequency-dependent I/O phase relationships for depolarization vs hyperpolarization. (left: V_memb_, right: Φ^*±*^(*f*); Φ^−^(*f*) depolarizing: dashed lines, Φ^−^(*f*) hyperpolarizing: solid lines; Φ^*±*^(*f*)*>*0 means V_memb_ peak/trough *leads* (precedes) stimulus peak/trough; Φ^*±*^(*f*) = 0 means V_memb_ peak/trough and stimulus peak/trough were synchronous; Φ^*±*^(*f*)*<*0 means V_memb_ peak/trough *lags* (follows) stimulus peak/trough). A. Small amplitude stimulus produced characteristic near-linear response (Φ^+^(*f*) and Φ^−^(*f*) overlaid in gray). B. Higher amplitude subthreshold stimulus showed mildly nonlinear amplitude response (peak 5% larger than trough) and differences in phase response (black line reproduced from A in B-D for reference). Resonance frequencies for depolarization and hyperpolarization were similar (6.4 and 6.6 Hz, respectively; Figure 2-1). C. Suprathreshold chirp with blocked spikes (simulated TTX experiment). Amplitude and phase differences are more pronounced than in B. D. Spiking patterns (dashed line) showed still greater phase lead. Hyperpolarizing responses were similar to those in C.

As previously demonstrated in L5b pyramidal neurons (Hutcheon et al., 1996; Ulrich, 2002; Dembrow et al., 2015) and this model neuron (Kelley et al., 2021), small, subthreshold chirp stimuli produced near-linear responses (Fig. 2A). For a stimulus amplitude of 0.025 nA, Φ^−^(*f*) and Φ^+^(*f*) were nearly identical to traditional impedance phase Φ(*f*). *I*_h_ produced “phenomenological inductance”, where the neuronal membrane, rather than behaving like a low-pass (RC), produced resonance and other properties resembling a band-pass (RLC) circuit without employing an actual inductor. Due to the phenomenological inductance generated by *I*_h_, input and output were in phase at the *synchrony point* (Φ = 0; here at 4.5 Hz), and there was inductive phase or phase lead (Φ *>* 0; output leads input) at frequencies below the synchrony point. There was phase lag (Φ *<* 0; output lags behind input) at frequencies higher than the synchrony point (Fig. 2A). For higher amplitude stimuli producing nonlinear V_memb_ response (maximum peak 5% larger than minimum trough), we analyzed the upper and lower envelopes of V_memb_ separately since they showed different frequency-dependent I/O phase relationships (Φ^+^(*f*) and Φ^−^(*f*), respectively) (Fig. 2B-D). Compared to the low-amplitude responses (Fig. 2B black line, copied from Fig. 2A in each panel), high-amplitude subthreshold stimuli produced less phase lead and a lower synchrony point (3 Hz) during depolarizing half-cycles (dashed blue), but they produced more phase lead and a higher synchrony point (6 Hz) during hyperpolarizing half-cycles (solid blue). We also computed an analog to impedance amplitude for the depolarizing and hyperpolarizing half-cycles developed by Pena et al. (2019) and found that, in contrast, the resonance frequencies for the two were nearly indistinguishable (6.4 and 6.6 Hz, respectively; Figure 2-1). Thus, small nonlinearities differentially altered the time course of depolarization and hyperpolarization with minimal change in frequency selectivity. For suprathreshold stimuli with spiking blocked, simulating a tetrodotoxin (TTX) experiment, both depolarizing and hyperpolarizing responses showed more phase lead than seen with either subthreshold example (Fig. 2C; note broadened y-axis); synchrony points were both shifted to higher frequencies: 5.5, 7.5 Hz, respectively.

As expected, spiking provided a major change in phase relationships for the depolarizations, with little change in phase relationships for hyperpolarizations (Fig. 2D). The initial burst and subsequent spikes occurred during the upswings of the input wave where the voltage reached threshold. The spike is itself a rapid depolarization which easily beats the chirp to the peak, providing the large phase lead of almost 1 radian. This phase lead continued at higher frequencies up to the synchrony point of 7.5 Hz, higher than in the other cases. For stimulus frequencies above 8.3 Hz, spikes did not occur on every stimulus cycle, giving the varying phase shift over successive cycles, alternately preceding or following the stimulus peak (Fig. 2D, purple dashed line).

I/O phase relationships for both hyperpolarizing and depolarizing responses in the high-subthreshold regime were dependent on the phenomenological inductance produced by *I*_h_(Fig. 3). Blocking *I*_h_ (simulating ZD7288 application) eliminated inductive phase and synchrony (Fig. 3A). Additionally, blocking *I*_h_ nearly eliminated the differences between the I/O phase relationships for hyperpolarization and depolarization, while retaining differences in V_memb_ amplitude (peak V_memb_ response 5% higher than trough due to the effect of voltage-gated Na^+^ channels, *I*_Naf_). Stimulus location altered phase due to the higher *I*_h_ density in the dendrites compared to the soma (Fig. 3B). Stimulating in apical dendrite (260 *μ*m from the soma) produced increased phase lead and higher synchrony points for both depolarization and hyperpolarization compared to stimulating the soma. Because *I*_h_ activation is voltage-dependent, phase was altered by changes in resting membrane potential (RMP) and stimulus amplitude. RMP decrement by constant negative current stimulus brought the voltages closer to the *I*_h_ half-activation point, so that hyperpolarizing responses were increased at decreased RMP (−5 and -10 mV from baseline of -74 mV), and increased activation of *I*_h_ increased phenomenological inductance (Mauro, 1961; Mauro et al., 1970).

**Figure 3:**
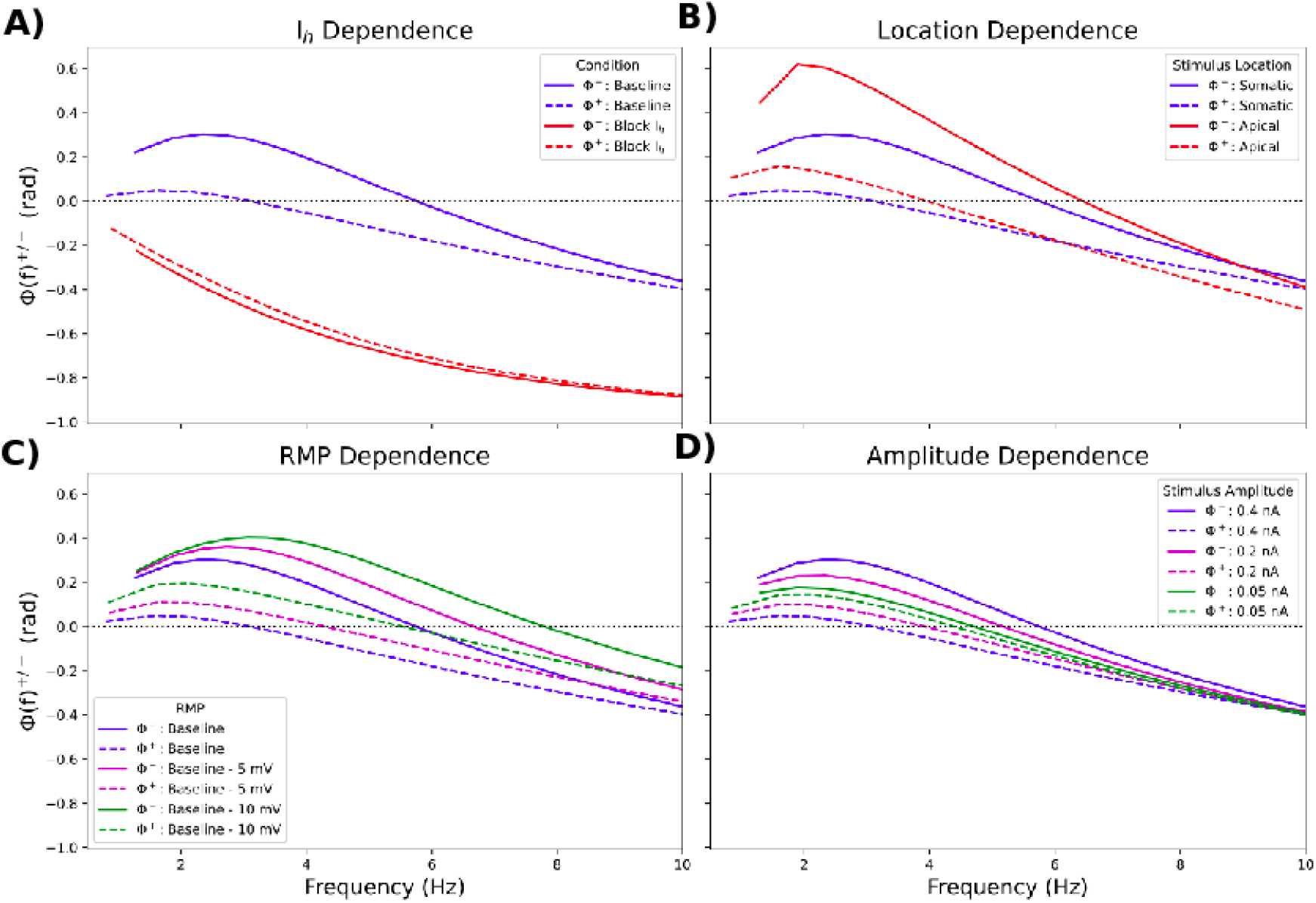
I/O phase shifts in subthreshold, nonlinear regime depended on *I*_h_ and stimulus parameters and location. Φ^+^(*f*) (dashed lines) and Φ^−^(*f*) (solid lines) A. when blocking *I*_h_ vs. control; B. when stimulating the apical dendrite (260 *μ*m from the soma) vs. the soma; C. under different artificially set RMPs; D. for different stimulus amplitudes.

Decreasing RMP by 10 mV increased phase lead and the synchrony points (by 2.1 Hz for hyperpolarization and 2.5 Hz for depolarization) (Fig. 3C). Changes in stimulus amplitude affected the degree to which different voltage-gated ion channels open (namely, HCN and voltage-gated Na^+^ channels), and thus the degree of nonlinearity in the V_memb_ response. Very small (0.05 nA) stimuli produced the near-linear response, giving negligible difference between the I/O phase relationships for depolarization and hyperpolarization (Fig. 3D green lines). With increasing stimulus amplitude, I/O phase relationships for hyperpolarization and depolarization diverge from the traditional impedance phase (Φ(*f*)): differences in the synchrony points for hyperpolarization and depolarization were caused by increasing *I*_h_ and *I*_Naf_, respectively (Fig. 3D).

We also examined phase shifts for spiking during a constant-frequency sinusoidal stimuli to determine how they changed with stimulus cycle, *n*. For particular combinations of stimulus amplitude and frequency, 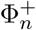 decreased over the course of multiple stimulus cycles, a phenomenon we termed *phase retreat* (Fig. 4A). Using an 8 Hz stimulus, the first spike (part of a burst) occurred before the peak of the first stimulus cycle (*n* = 0). 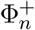 then decreased over subsequent cycles due to spike adaptation Spikes occurred after the stimulus peak from the fourth stimulus cycle onward (*n* = 4) until reaching a steady-state above *n* = 8 where 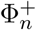 changed little. Thus, during phase retreat, 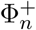 encoded the duration of the stimulus prior to reaching steady-state. Phase retreat was primarily due to *I*_h_ and *I*_AHP_ (SK channels) Fig. 4B), as *I*_h_ inactivated and *I*_AHP_ activated due to depolarization and subsequent Ca^2+^ entry (SK is a Ca^2+^-dependent K^+^ channel). Blocking *I*_h_ or *I*_AHP_ eliminated changes in 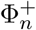 over multiple cycles, eliminating 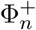 encoding of stimulus duration. The phase curve shifted downward (more delayed) with dendritic localization of stimulation due to the conductance delay from apical dendrite to soma; more subtle changes in phase shift across cycles could be seen which was due to channel density differences between dendrite and soma (Fig. 4C). Phase retreat was frequency dependent since low frequencies did not produce significant adaptation and in fact provided a slight facilitatory effect (Fig. 4D). Phase retreat was most pronounced at 8 Hz, the fastest following frequency for this stimulus amplitude. At 9 Hz the cell could not follow, producing dropped spikes with corresponding discontinuous phase relation. Increasing stimulus amplitudes produced less phase retreat as threshold on each cycle was reached before the current peaked (Fig. 4E).

**Figure 4:**
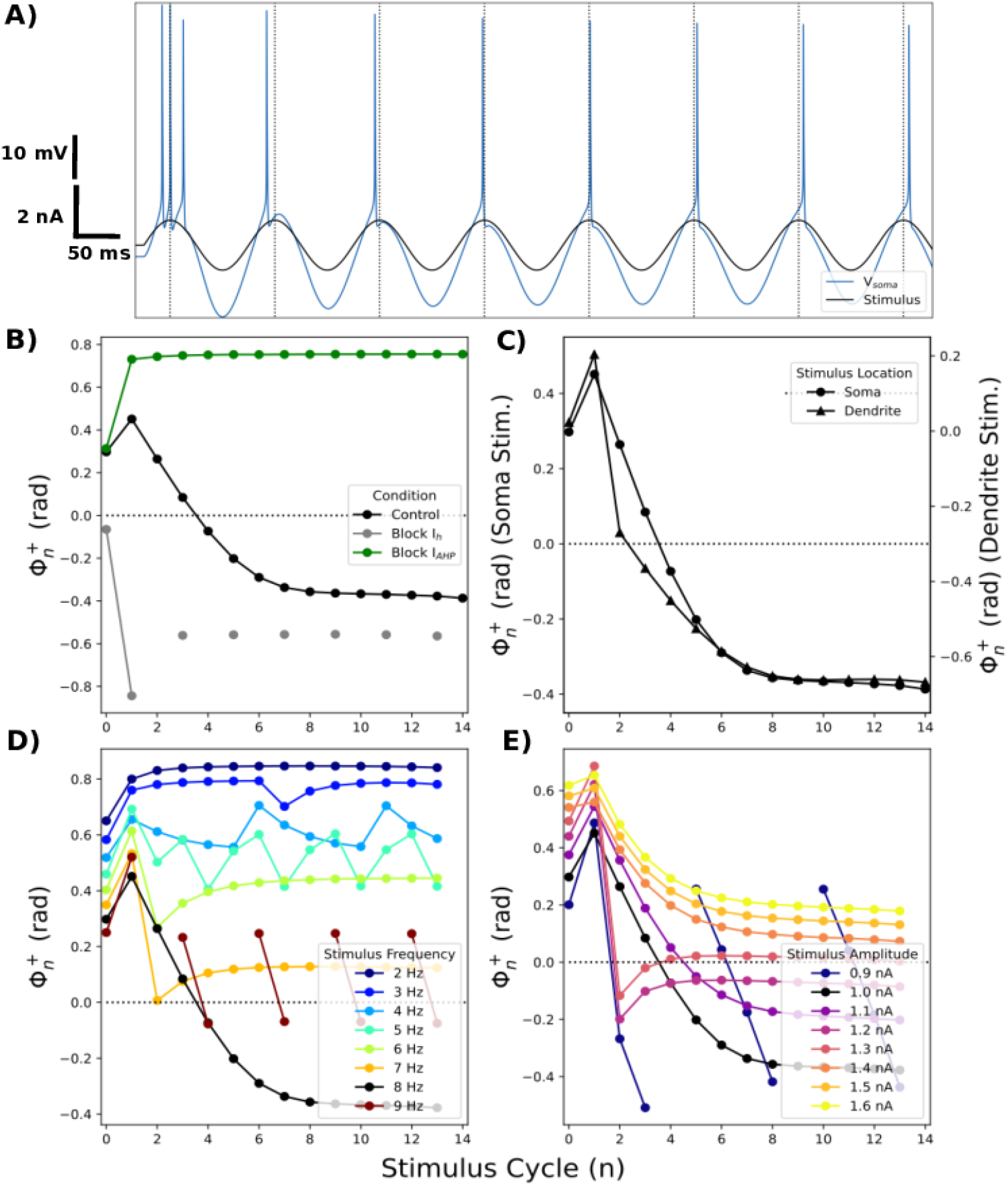
Sinusoidal stimuli produced phase retreat. A. 8 Hz sinusoidal current stimuli (black) was injected into the soma. Resulting changes in V_memb_ were superimposed (blue). Spikes occurred later on each stimulus cycle. Note changing distance between spikes and 0 radian stimulus phase (vertical dashed line) between cycles. We called this pattern phase retreat. B. Phase retreat was eliminated by blocking *I*_h_ (gray) or *I*_AHP_ (green). C. Phase retreat was location dependent. Stimulus amplitude was increased for dendritic stimulation to get similar spiking at soma. Note separate y-axes for dendritic and somatic stimulation. D. Phase retreat was frequency dependent. Resulting 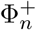 for spikes from stimuli of varying frequencies and amplitude of 1 nA. E. Phase retreat was amplitude dependent. Resulting 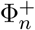 for spikes from 8 Hz stimuli of varying amplitudes.

Since the spiking response to sinusoidal stimulus was nonstationary, we examined which if any phase relation was predicted by the chirp stimulus (Fig. 5). Prediction was strong for the steady-state (adapted) depolarization phase (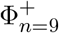 compared to chirp Φ^+^(*f*)), while the phase shift during earlier stimulus cycles 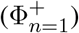 was not captured except at low frequencies where little change in phase took place (Fig. 4D).

**Figure 5:**
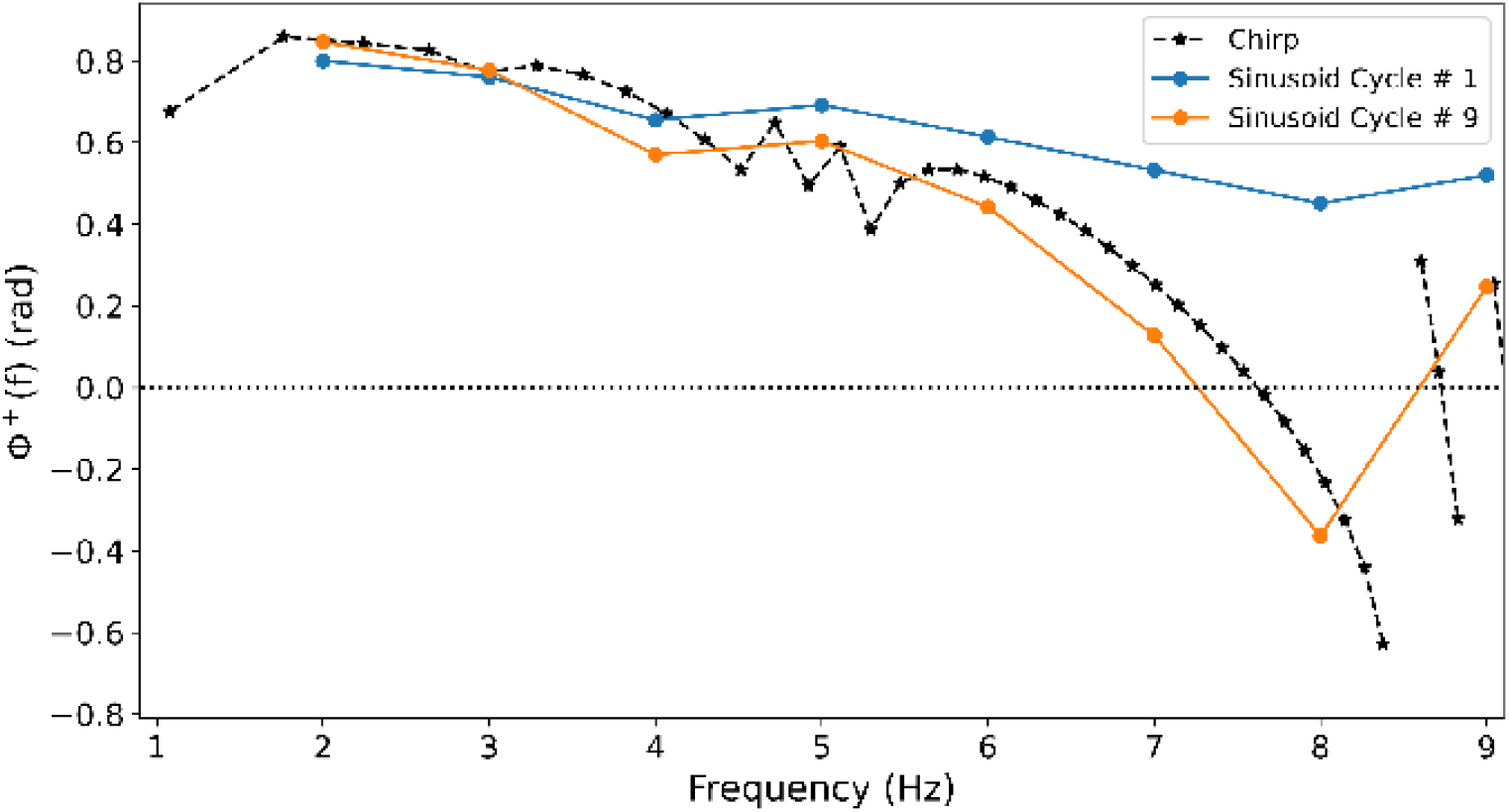
Suprathreshold chirp stimulation predicted steady-state 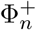. 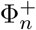 for n=1 (blue), n=9 (orange) compared to Φ^+^(*f*) from chirp (black, reproduced from Fig. 2D).

We simulated synaptic/input facilitation by linearly increasing amplitude during the sinusoidal stimuli (Fig. 6). As expected, 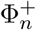 increased over the course of multiple stimulus cycles as the cell reached spike threshold earlier in the stimulus cycle than on the previous. Stimulating the soma with an 8 Hz augmenting sinusoid (initial peak: 0.11 nA; increase: 0.04 nA/cycle) produced *phase advance*: 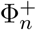 increasing on each cycle, with phase lag on first cycle and phase lead on later cycles (Fig. 6A). Similar to phase retreat, during phase advance 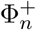 encoded the amplitude and duration of the stimulus prior to reaching steady-state. Strong encoding of stimulus amplitude/duration only lasted for five stimulus cycles before 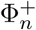 reached steady state in baseline conditions (Fig. 6A). Phase advance was modulated in opposing directions by *I*_h_ vs *I*_AHP_ blockade (Fig. 6B). Blocking *I*_AHP_ enhanced 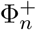 encoding of stimulus amplitude/duration by increasing range of 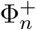 and increasing 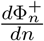 for more stimulus cycles. However, there was spike failure on cycles 11–13 (Fig. 6B, interruption in green curve). Blocking *I*_h_ also augmented 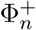 encoding of stimulus amplitude/duration, increasing 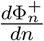 for more stimulus cycles. Phase advance was location dependent (Fig. 6C – note elimination of delay due to location), with increased 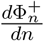 produced by increased influence of ion channels in apical dendrite. Phase advance was strongly frequency dependent (Fig. 6D). Rate of stimulus amplitude change had relatively little influence on Φ^+^, providing consistency of phase encoding of stimulus duration (Fig. 6E).

**Figure 6:**
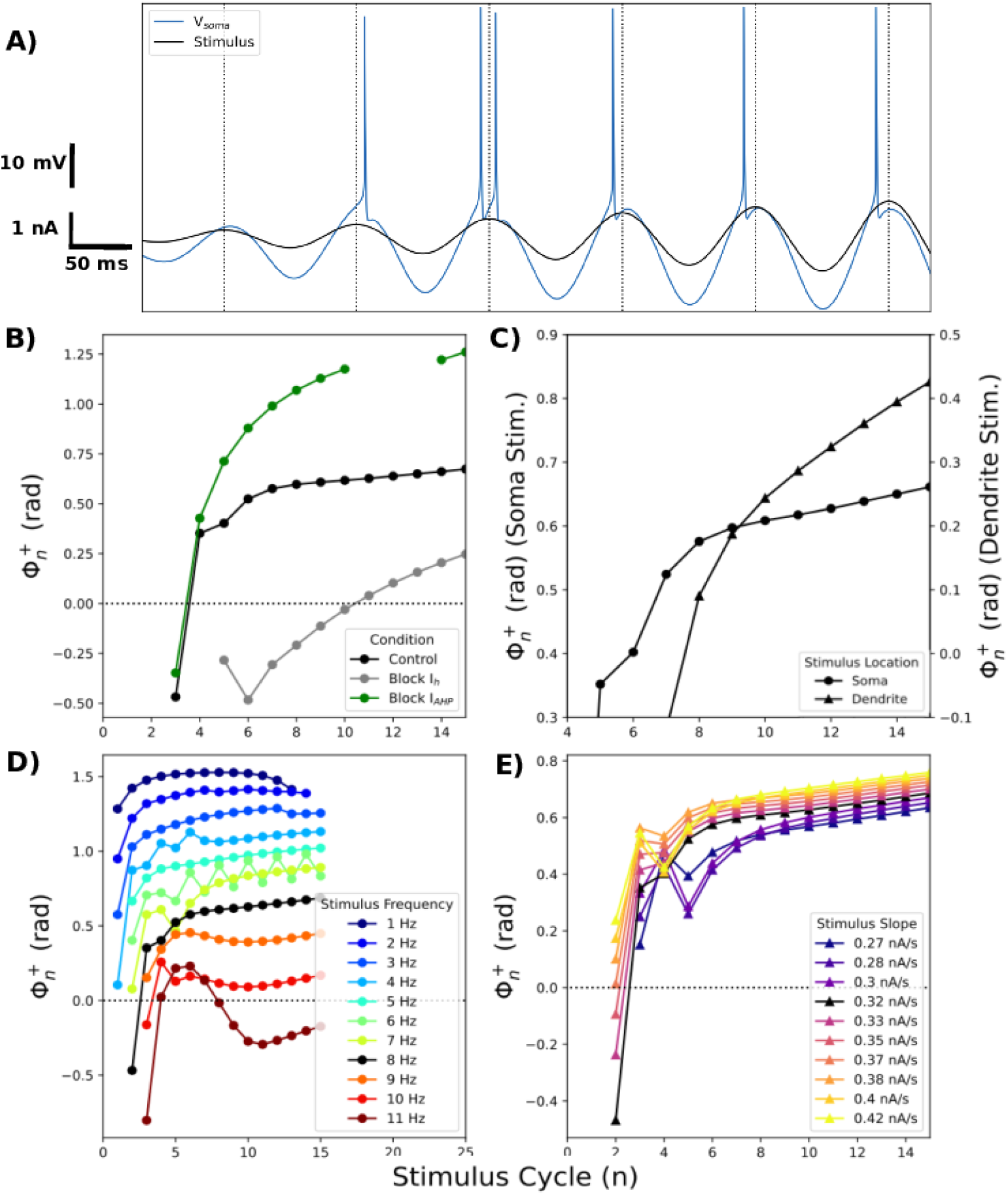
Increasing stimulus amplitude over time produced phase advance. A. 8 Hz sinusoidal current stimuli whose amplitude increased 0.04 nA/cycle (black) was injected into the soma. Resulting changes in V_memb_ were superimposed (blue). Spikes occur progressively earlier on each stimulus cycle. Note changing distance between spikes and 0 radian stimulus phase (vertical dashed line) between cycles. We called this pattern phase advance. Stimulus amplitude was 0.11 nA for first peak shown. B. Blocking *I*_h_ and *I*_AHP_ modulated phase advance. C. Location dependence: phase advance was stronger when stimulating the apical dendrite (260 *μ*m from than the soma); separate y-axes for dendritic vs somatic stimulation to remove phase delay due to distance from soma recording. D. Phase advance was frequency dependent. 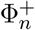 for spikes elicited by stimuli of different frequencies whose amplitude increased 0.32 nA/s E. Rate of stimulus amplitude increase had little effect on phase advance. 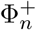 for spikes elicited by 8 Hz stimuli whose amplitude increased at various rates. Note differences in axes across panels.

## Discussion

Classical impedance analysis in neurophysiology requires low-amplitude signals in order to remain close to a linear ideal: the resistor-inductor-capacitor (RLC) circuit (Fig. 2A) (Kelley et al., 2021; Cole, 1941; Mauro, 1961). These small signals only slightly activate voltage-sensitive ion channels, giving rise to a phenomenological inductance which can then be further analyzed using traditional electrical engineering techniques. However, small-signal inputs do not help us understand the strongly nonlinear responses from the far higher amplitude signals characteristic of neural inputs. In this paper, we identified cellular I/O phase relationships under depolarization and hyperpolarization to sinusoidal and chirp inputs of different stimulus amplitudes and frequencies; under different membrane conditions (blockade of specific membrane conductances); in subthreshold, spike-blocked suprathreshold, and spiking regimes. These results are important for understanding the response of neurons to the incoming oscillatory inputs that create the field potential oscillations identified as delta – beta waves, suggesting different phases for responses to inhibitory vs. excitatory postsynaptic potentials.

We defined a *synchrony point* as the cross-over frequency from phase lead to phase lag where input and output are in phase (Φ^*±*^ = 0). Classical low-amplitude signal responses produced very similar phase relationships for hyperpolarizing and depolarizing half-cycles, sharing a single synchrony point (Fig. 2A). By contrast, higher amplitude subthreshold V_memb_ responses showed more phase lead and a higher synchrony point for hyperpolarization compared to depolarization due to increased *I*_h_ (Fig. 2B). Phase lead and synchrony were dependent on the phenomenological inductance generated by *I*_h_ and its interactions with different voltage-gated ion channels activated during stimulation (Fig. 3). Blocking *I*_h_ eliminated the phenomenological inductance and effectively abolished the differences between the I/O phase relationships for depolarization and hyperpolarization, without however abolishing the asymmetry of the V_memb_ amplitude in the high-signal subthreshold domain (Fig. 3A). Phenomenological inductance serves to compensate for delays in V_memb_ changes caused by membrane capacitance (Kelley et al., 2021; Vaidya and Johnston, 2013), and our results show that this compensation will be augmented for large inhibitory inputs and diminished for large excitatory inputs compared to smaller amplitude inputs that remain in the linear regime. Vaidya and Johnston (2013) posited that the phenomenological inductance generated by gradients of *I*_h_ along the dendrites ensure synchronous synaptic stimuli are simultaneously integrated at the soma, regardless of the synapse location – a “democracy of synapses.” Our results suggest that when stimuli are large, causing V_memb_ to approach spike threshold, the representation of excitatory and inhibitory stimuli becomes undemocratic.

Although Φ^+^ changed smoothly with suprathreshold chirp stimulus during spike blockade (Fig. 2C), chirp stimulation was of limited utility for characterizing phase relationships with spiking (Fig. 2D, Fig. 5). This shortcoming was due to the combination of cell nonstationarity with the nonstationary input. The I/O phase relationships for spiking identified by chirp predicted only the steady-state 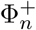 (Fig. 5). For an 8 Hz stimulus, steady-state 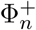 is not achieved for nearly 1 second, a long time in terms of brain information processing.

We identified two patterns of spiking in response to continuing sinusoidal stimuli: **1**. *phase retreat*, in which spikes or bursts occurred progressively later in the cycles of the input stimulus (Fig. 4), and **2**. *phase advance*, in which spikes or bursts occurred progressively earlier in the cycles of the input stimulus (Fig. 4). Phase retreat emerged from spike-frequency adaptation – *I*_h_ and *I*_AHP_ were required to produce the gradual increase in spike threshold, driving phase shift 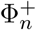 lower until reaching steady-state. Phase advance, on the other hand, emerged as the result of a sinusoidal stimulus whose amplitude increased over time, with the phase shift 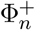 increasing over time. Phase advance was also strongly modulated by, but not dependent on, *I*_h_ and *I*_AHP_. Both phenomena were also affected by stimulus location, frequency, and amplitude. Phase advance and phase retreat represent modes of spiking activity in which rate coding and temporal coding coexist. In both cases, information about the stimulus frequency is encoded in the spike rate. Information about the stimulus duration is encoded by 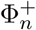 during phase retreat, and information about the stimulus amplitude, whether due to synaptic potentiation, presynaptic population recruitment, or increased presynaptic firing rate, is encoded by 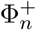 during phase advance.

Phase shift relative to the local field potential (LFP) suggests phase coding as an adjunct to rate coding in the central nervous system (CNS) (Magee, 2001; Mehta et al., 2002; Harris et al., 2002; Fries et al., 2007; Leung, 2011). CNS phase coding has been most thoroughly studied in the context of *phase precession* in hippocampal place cells, but has also been seen in entorhinal grid cells(Hafting et al., 2008), medial prefrontal cortex (Jones and Wilson, 2005), and ventral striatum (van der Meer and Redish, 2011). Large pyramidal neurons in all of these areas have much in common in their impedance profiles, expression of HCN channels, and morphological features (Das et al., 2017; Hutcheon et al., 1996; Ulrich, 2002; Leung and Yu, 1998). In phase precession, spiking occurs earlier on successive cycles of the theta wave as an animal moves towards the center of a place field (O’Keefe and Recce, 1993). By contrast, a recent study identified “phase roll” in hippocampal CA1 neurons, with phase moving in the other direction to later times relative to theta (Sloin et al., 2022).

While phase precession has frequently been modeled as an emergent property of the network (Geisler et al., 2007; Jensen and Lisman, 1996; Bose and Recce, 2001; Wallenstein and Hasselmo, 1997; Tsodyks et al., 1996), we suggest here that intrinsic cell properties would also play a role, complementing network properties. In this view, phase advance would increase phase precession with increased sensory input, while phase retreat would produce phase roll in other cells, with steady inputs during the same theta period. Network action would augment intrinsic effects by complementing the oscillatory excitatory stimuli in dendrites with inhibitory stimuli in the soma (Kamondi et al., 1998; Magee, 2001; Mehta et al., 2002; Leung, 2011). Our results predict that phase precession could be tuned through modulation of *I*_h_ or *I*_AHP_ (Fig. 6B). Our model predicts that intrinsic aspects of both phase precession and phase roll could be tested by intracellularly applied blockers (intracellular to avoid network effects): precession would be enhanced by blocking either *I*_h_ or *I*_AHP_; roll would be eliminated by either.

The hypothesis of coexisting causes for phase precession comes as no surprise in the complex multiscale dynamical systems of the brain. The two hypotheses – network cause or single-cell cause – are complementary. Hence there is no falsifiability through eliminating one from the system as there would be with a physics hypothesis. Furthermore, eliminating intrinsic effects on a large scale (not just a single cell) or eliminating network effects will produce a new dynamical system that will not speak to either hypothesis: *e*.*g*., eliminating the network through block of all synaptic connections (locally, so as not to kill the animal), will eliminate precession trivially. The difficulties of multiscale modeling, and multiscale concepts, in biology are seen to be of a different order than those in many fields because of the failure of scale-by-scale encapsulation. Famously, the individual atoms of a gas can be encapsulated as particles for the purposes of understanding the gas laws and thermodynamics: there is no need to consider electron shells or intranuclear forces. By contrast, the scale of a particular ion channel type (*e*.*g*., *I*_h_) cannot be fully encapsulated as a phenomenological inductor for the purpose of understanding neuronal membrane dynamics; the scale of the single neuron cannot be fully encapsulated as a sum-and-squash point neuron for understanding network dynamics; the scale of thalamus cannot be encapsulated as part of thalamocortical dynamics; *etc*. This failure of encapsulation (a complication of scale-overlap) is seen throughout biology and manifests particularly clearly in brain. Understanding neural systems, and the complex behaviors they give rise to, involves interactions across spatiotemporal scales.

## Acknowledgements

We wish to thank Dr. Michele Migliore (National Research Council of Italy), Dr. Salvador Dura-Bernal (SUNY Downstate), Dr. Michael Hasselmo (Boston University), and Dr. Steven Fox for useful discussions about this project. This research was funded in part by Aligning Science Across Parkinson’s [ASAP-020572; WWL] through the Michael J. Fox Foundation for Parkinson’s Research (MJFF). For the purpose of open access, the author has applied a CC BY public copyright license to all Author Accepted Manuscripts arising from this submission.

## Extended Data

**Figure 2-1.**
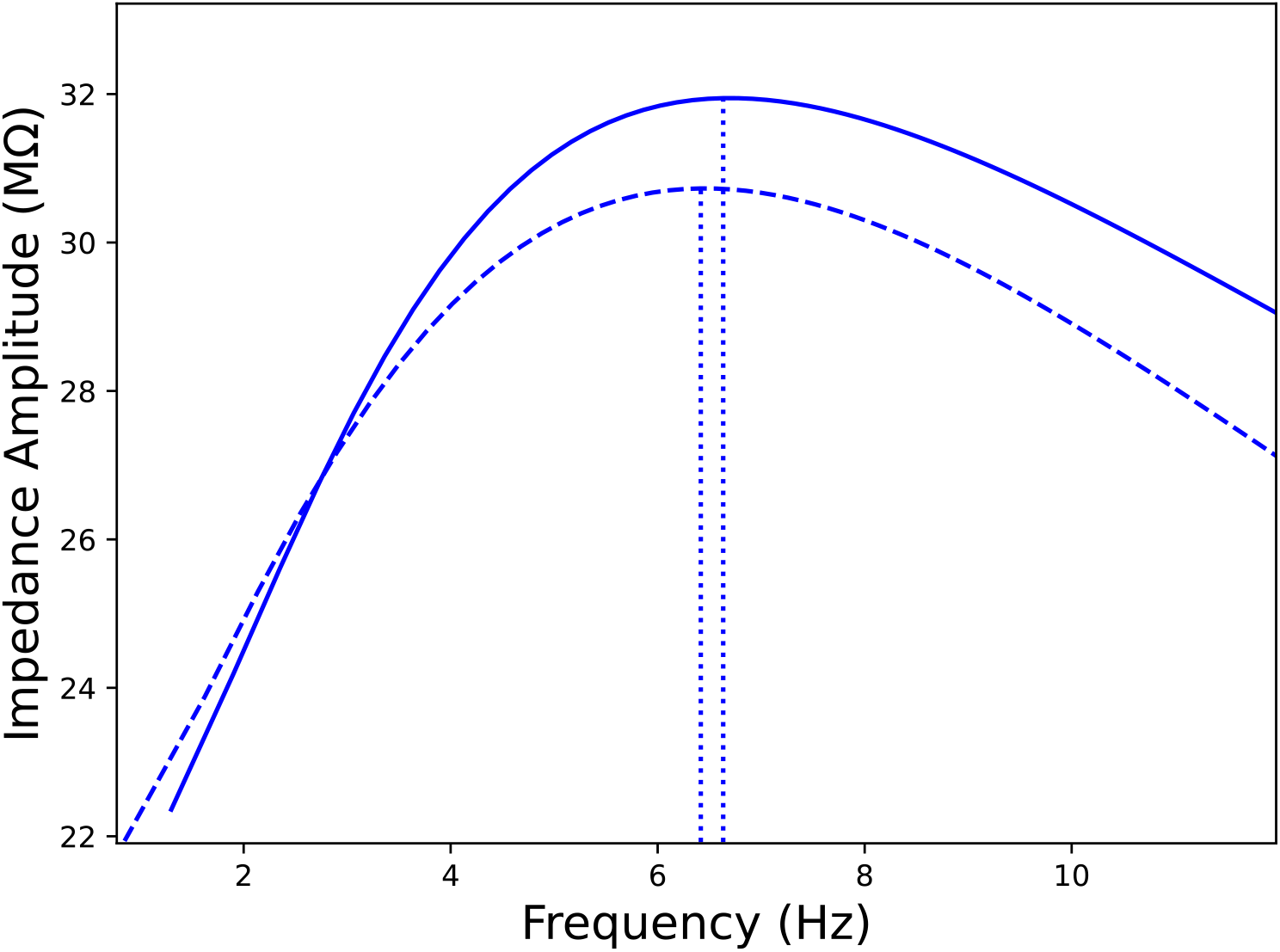
Nonlinear, subthreshold V_memb_ response exhibited similar resonance frequencies during depolarization and hyperpolarization. Nonlinear analogs of impedance amplitude (as described in Pena et al. 2019) for depolarizing (dashed) and hyperpolarizing (solid) half cycles of V_memb_ response to 0.4 nA amplitude stimulus (see Fig. 2A for V_memb_ trace). Resonance frequencies are indicated by vertical dotted lines.

